# Plant sedimentary ancient DNA from Far East Russia covering the last 28 ka reveals different assembly rules in cold and warm climates

**DOI:** 10.1101/2020.12.11.406108

**Authors:** Sichao Huang, Kathleen R. Stoof-Leichsenring, Sisi Liu, Jeremy Courtin, Andrej A. Andreev, Luidmila. A. Pestryakova, Ulrike Herzschuh

## Abstract

Woody plants are expanding into the Arctic under a warming climate. The related impact on plant diversity is not well understood because we have only limited knowledge about plant assembly rules and because of a lack of time-series of plant diversity. Here, we applied sedimentary ancient DNA metabarcoding using the plant-specific *g* and *h* primers of the *trnL* gene to a sediment record from Lake Ilirney (central Chukotka Far Eastern Russia) covering the last 28 thousand years. Our results show that forb-dominated steppe-tundra and Saliceae-rich dwarf-shrub tundra communities dominated during the cold climate before 14 ka, while deciduous erect-shrub tundra was abundant during the warm period between 14 and 0 ka. *Larix* invasion during the late Holocene substantially lagged behind the period of densest vegetation and likely warmest period between 10 and 6 ka. Overall, we discovered the highest richness during 28–23 ka and a second richness peak during 13–10 ka: both periods are characterised by low shrub abundance. The richest communities during the cold pre-14 ka period were phylogenetically clustered, which probably originates from environmental filtering along with niche differentiation under limited resources. In contrast, the richest post-14 ka community was phylogenetically overdispersed, likely originating from an erratic recruitment process in the course of warming. Despite differences in timescale, some of our evidence can be relevant to arctic plant diversity changes. By analogy to the past, we expect a lagged response of tree invasion. In the long-term, ongoing expansion of deciduous shrubs will eventually result in a phylogenetically more diverse community but will also cause reduced plant taxonomic richness; however, richness may overshoot in the short-term.

## Introduction

Recent warming has triggered vegetation changes (Wilson and Nilsson 2009). For example, woody taxa have expanded their range further north (Kharuk et al. 2006, Pearson et al. 2013) albeit at a rather low rate compared to the corresponding temperature range (Kruse et al. 2018). Arctic plant richness is considered sensitive to climate change (Callaghan et al. 2004) because plant richness of tundra is often lower than in boreal forests (Khitun et al. 2016). Some modern observations suggest an increased diversity in arctic tundra related to forest expansion under a warming climate (Rupp et al. 2001, Danby et al. 2011), although other studies suggest a decline (Walker et al. 2006). Change in phylogenetic diversity is of interest as it can indicate changes in assembly rules under warmer and colder climate conditions. Low phylogenetic diversity reflects the co-occurrence of closely related species and suggests habitat filtering. In contrast, a community with high phylogenetic diversity hosts an assembly of distantly related species (Webb 2000, Webb and Pitman 2002). On the one hand, phylogenetically more varied plant species have been found to coexist, when, as a result of warming, more soil nutrients are available (Chu and Grogan 2010, Zhu et al. 2020). On the other hand, climate warming may decrease phylogenetic diversity (Thuiller et al. 2011) through environmental filtering due to drought (Dorji et al. 2012). Accordingly, the major rules of the plant assembly process under the ongoing warming are still not well understood and such predictions about future vegetation composition as well as taxonomic and phylogenetic diversity are uncertain.

The knowledge gap with respect to assembly rules in plant communities largely originates from a lack of long-term studies in remote arctic areas. Accordingly, proxy information such as plant records from lake sediments needs to be explored. Conventional studies of vegetation history are mostly based on pollen analysis (Moser and MacDonald 1990, Anderson and Brubaker 1994, Seppä et al. 2002, Andreev et al. 2011). Many Eurasian pollen records show lower palynological richness during the cold and dry Late Glacial interval compared with the warm and wet Holocene (Feurdean et al. 2011, Chytrý et al. 2017), although there are studies that indicate a relatively high pollen-taxa diversity during the Late Glacial (Blarquez et al. 2014, Reitalu et al. 2015) and a decreasing diversity during the Holocene (Blarquez et al. 2014). Sedimentary pollen records from north-eastern Europe show that the climate conditions led to an increase in overall plant richness but a decrease in phylogenetic diversity during the post-glacial period (Reitalu et al. 2015). Hence, our knowledge on past changes in phylogenetic alpha diversity is still very scarce, especially for Siberia, and thus our information about major plant assembly rules in times of warming is insufficient.

Knowledge gaps about millennial-scale composition and diversity changes may, to some extent, arise from the limited taxonomic resolution and taphonomic peculiarities of pollen records. For example, *Larix*, the treeline-forming genus in most of Siberia is poorly reflected in pollen records (Sjögren et al. 2008, Binney et al. 2011, Niemeyer et al. 2017). Plant DNA preserved in lake sediments, however, can provide a reliable proxy for vegetation composition and diversity (Bálint et al. 2018). Modern studies have confirmed that sedimentary DNA from high-latitude regions reliably reflect the vegetation surrounding the lake (Parducci et al. 2019) and can be used to trace temporal changes in plant diversity (Niemeyer et al. 2017, Zimmermann et al. 2017). The universal, plant-specific and short barcode marker of the P6 loop of the chloroplast is used for sedimentary DNA (sedaDNA) studies (Taberlet et al. 2007).

Plant DNA metabarcoding has revealed that Siberian plant diversity was highest during pre-LGM (i.e. Last Glacial Maximum about 21 ka) and the lowest during the LGM (Willerslev et al. 2014, Epp et al. 2018, Clarke et al. 2019). This was speculatively related to changes in forb abundance and the impact of megafauna (Willerslev et al. 2014) but Clarke et al. (2019) hypothesised that the growth of shrub taxa during the Late Glacial led to competition within arctic taxa, resulting in reduced species diversity.

Our study investigates whether plant assembly rules vary by assessing changes in vegetation composition and taxonomic and phylogenetic alpha diversity as inferred from sedimentary ancient DNA from Lake Ilirney (Chukotka, Far East Russia) since 28 ka. We aim (1) to reconstruct vegetation composition changes; (2) to investigate changes in taxonomic alpha diversity (richness, evenness) and its relationship with vegetation compositional changes, particularly during woody taxa invasion; and (3) to explore changes in phylogenetic diversity and how this reflects variations in vegetation composition and taxonomic diversity.

## Methods

### Study area

Lake Ilirney (67°21’N, 168°19’E) is a glacially formed lake located in the autonomous region of Chukotka (Far East Russia, Fig. 1). The lake is bounded by the Anadyr Mountains (up to 1790 m a.s.l.) to the north. According to the meteorological station at Ilirney, mean annual temperature is −13.5°C, and mean January and July temperatures are −33.4 and 12.1 °C, respectively. Annual precipitation is ~200 mm (Menne et al. 2012).

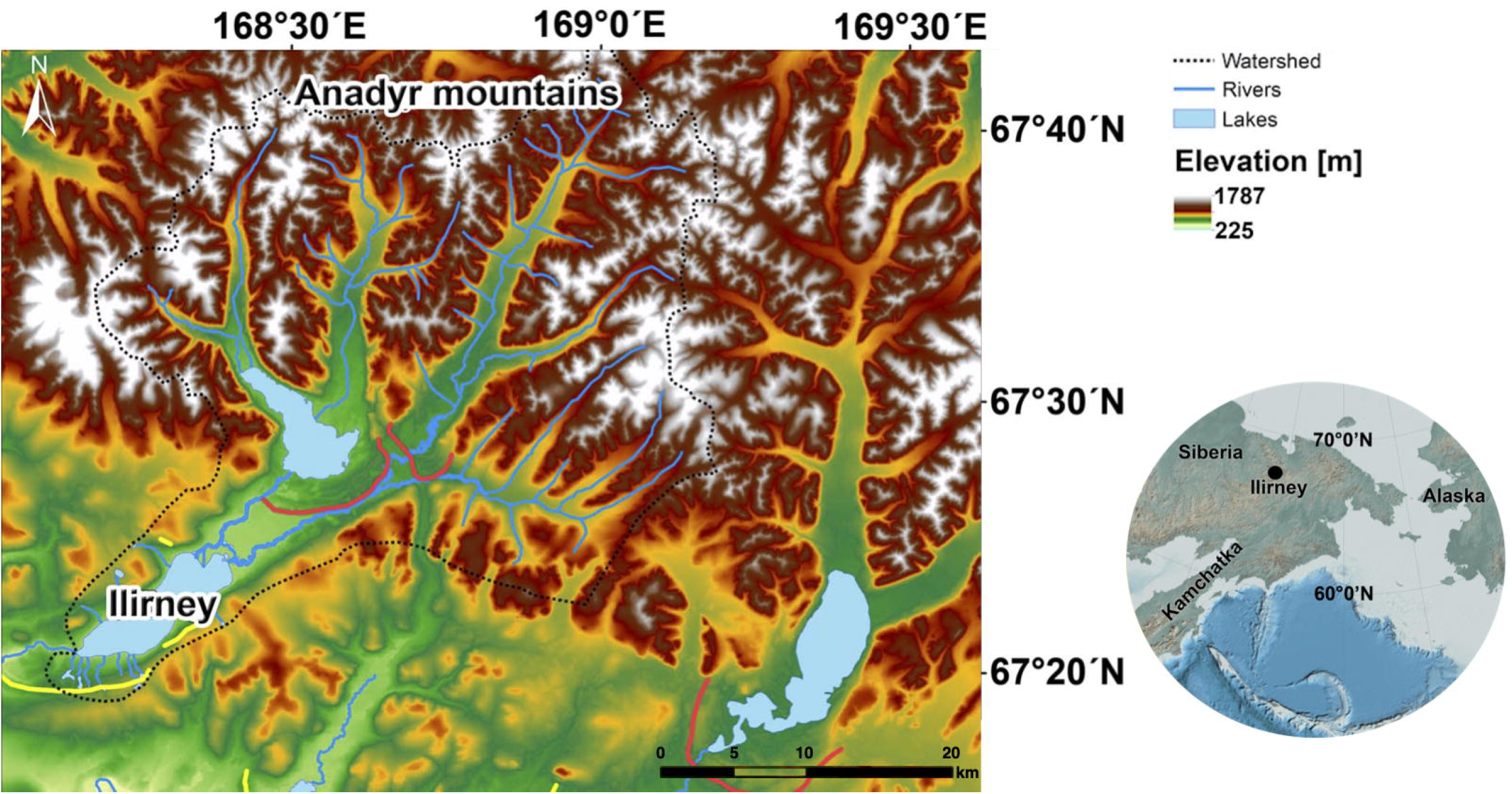

Lake Ilirney is situated at the tundra-taiga ecotone, which is characterised by a mixture of forest tundra, shrub tundra, and open tundra. Stands of *Larix cajanderi* are common in the lake vicinity as well as low and dwarf shrubs such as *Salix* spp., *Betula nana, Ledum palustre, Vaccinium* spp., and *Empetrum nigrum.* Elevational transitions between forest tundra and open tundra are often covered by patchy dwarf pine *(Pinus pumila).* Open tundra on the gentle slopes (540–700 m a.s.l) is graminoid-rich including hummock (*Eriophorum vaginatum*) and non-hummock *(Carex* spp.) tundra with dense moss cover in the ground layer. Intermediate elevations (700–900 m a.s.l.) are dominated by *Dryas octopetala* and lichens, accompanied by herbs from the Fabaceae, Orobanchaceae, Poaceae, Rosaceae, and Saxifragaceae. The vegetation at higher elevations becomes barren (Shevtsova et al. 2020).

### Sampling and dating

A 235 cm long sediment core 16-KP-01-L02 Long 3 was obtained from Lake Ilirney at 67.34148°N, 168.30443°E. Transportation, preservation and subsampling of the core for ancient DNA samples are described in detail in Appendix 1 in the Supplementary Material. The age-depth model is based on Accelerator Mass Spectrometry (AMS) radiocarbon dating of seven bulk total organic carbon samples from this core (Andreev et al. 2020 submitted) and correlation to a nearby 1040 cm sediment core with 25 dates (Vyse et al. 2020). Depths and the according ages of the samples used in this study are available in Pangaea *(DOI will be provided after the acceptance of the manuscript*).

### Genetic laboratory works

In total, 58 core sediment samples were selected for DNA extraction ranging from 1 to 235 cm depth. Approximately 3 g of sediment per sample was used for DNA extraction. The extraction and purification steps are detailed in Appendix 2 in the Supplementary Material. Briefly, DNA extracts were amplified using the plant-specific *g* and *h* primers of the *trnL* gene (Taberlet et al. 2007). Polymerase chain reaction (PCR) was performed on three replicates, followed by purification, pooling, and finally sequencing on the Illumina platform (Supplementary Material Appendix 2).

### Processing the sequence data

Sequence data were processed using the OBITools package (Boyer et al. 2016) by following the pipelines outlined in Appendix 3 of the Supplementary Material. The reference used for taxonomic assignment is based on the curated arctic and boreal vascular plant and bryophytes database (Sønstebø et al. 2010, Willerslev et al. 2014, Soininen et al. 2015). The data were further filtered as described in Appendix 3 of the Supplementary Material.

To obtain normalised count data, the true counts were resampled 100 times based on the minimum number of counts (84,343) across the samples (https://github.com/StefanKruse/R_Rarefaction). The final dataset for statistical analysis used the mean values of the 100 resampled datasets.

### Statistical analyses

Statistical analyses were computed in R 3.6.0 (R Core Team 2017). A double-square root followed by a Chord transformation (Legendre and Borcard 2018) was applied to the community compositional data prior to all statistical analyses. Analyses were performed in the “vegan” (Oksanen et al. 2013) and “rioja” packages (Juggins 2009). Relative count proportions of taxa within a sample were calculated to visualise taxa abundance for all ages. The transformed proportions were converted to Euclidean dissimilarity distances by *vegdist*. Hierarchical clustering was CONISS constrained using the function *chclust*. Zonation was analysed by the broken-stick model using *bstick.* Principal component analysis (PCA) was restricted to those taxa that occur at a minimum of 0.2% and in at least three samples.

Richness is presented as the mean value of the 100 resampled datasets. The exponential of Shannon’s index and the inverse of Simpson’s index were calculated based on the mean values of the resampling repeats using the function *diversity* in “vegan” (Oksanen et al. 2013). Evenness was calculated as the exponential of Shannon’s index divided by log(richness). Error bars are the minimum and maximum values of the resampled data. The taxonomic uniqueness of the samples and the taxa that contribute the most to these compositional differences were evaluated by local contribution to beta diversity (LCBD) and species contribution to beta diversity (SCBD). They were calculated using *beta.div* and the Euclidean method in the “adespatial” package (Dray et al. 2017) in both abundance- and occurrence-based communities.

The phylogenetic tree was produced based on implemented mega-trees of vascular plants using the function *phylo.maker* with the “scenario 1” approach in the “V.Phylomaker” package (Jin and Qian 2019). The following phylogenetic analyses were performed in the “picante” package (Kembel et al. 2010). We supplied the phylogenetic tree with the community data using *match*.*phylo*.*comm*. The cophenetic distance was calculated using *cophenetic*. The net relatedness index (NRI) and nearest taxon index (NTI) were used to quantify the degree of phylogenetic relatedness (Webb 2000). We created 999 null communities by randomly assembling communities using an “independent swap” algorithm. The null communities were used to compare the standardised effect sizes of mean phylogenetic distance (MPD) and mean nearest phylogenetic taxon distance (MNTD) to their observed communities implemented with *ses.mpd* and *ses.mntd,* respectively. We obtained the NRI by multiplying the estimates of MPD (mpd.obs.z) by −1 and NTI by multiplying the estimate of MNTD (mntd.obs.z) by −1.

We calculated the inter-community mean pairwise distance using the function *comdist*. The phylogenetic local contribution to beta diversity (pLCBD) was estimated using the function *LCBD.comp* in the *“* adespatial” package (Dray et al. 2017) for both abundance- and occurrence-based communities. The Pearson’s correlation between species composition, taxonomic, and phylogenetic diversity was calculated using *corr.test* in “psych” package (Revelle 2017).

## Results

### Overview of the sequencing data and taxonomic composition

Our raw sequencing data contain 45,103,727 paired reads and 12,816 sequence types. After stepwise filtering, the final dataset of 58 samples contains 21,697,725 reads and 158 sequence types with 100% identify to reference taxa. Twelve sequence types are identified to family including subfamily level, 79 sequence types are identified to genus or tribe and subtribe level, and 67 to species level. Aside from a few outliers, the number of reads per sample is of the same order of magnitude (median: 319,778). The plant community is dominated by shrubs and herbs. *Picea*, *Larix*, *Populus,* and *Ulmus* are the only tree taxa in our dataset and occur with low counts. 79.5% of the total counts are from shrubs and dwarf shrubs (Saliceae, *Alnus alnobetula, Dryas* sp.), and 20.4% are from herbaceous plants *(Eritrichium* sp., *Papaver* sp., Anthemideae).

According to the clustering and zonation results, the plant community can be divided into three main zones (Fig. 2). *Eritrichium* sp. is the most dominant taxon in Zone 1 (28–19 ka) having a mean relative abundance of 28.7%. Zone 1 features prevalent and diverse herbaceous plants. Abundant dwarf-shrubs and herbs include *Cassiope tetragona, Dryas* sp., *Papaver* sp., Anthemideae 1, and *Saxifraga* sp. 2. Saxifragaceae and Asteraceae are the most diverse families in zone 1, represented by 14 and 11 different sequence types, respectively. Saliceae and Betulaceae increase from Zone 1 to Zone 2 (18–14 ka) and show strong variation. There is an overall diversity reduction in Zone 2: the most diverse family, Saxifragaceae, contains only 7 sequence types. Zone 3 (14–0 ka) is dominated by woody taxa from Salicaceae, Betulaceae, and Ericaceae families. Saliceae is again the most abundant taxon. Another shrub *Alnus alnobetula* is the second dominant taxon with a high abundance between 9 and 6 ka in parallel to records of *Populus* and *Ribes*. There is a noticeable rise in the diversity of dwarf shrubs in Zone 3. Ericaceae is the most diverse family, represented by 13 different sequence types.

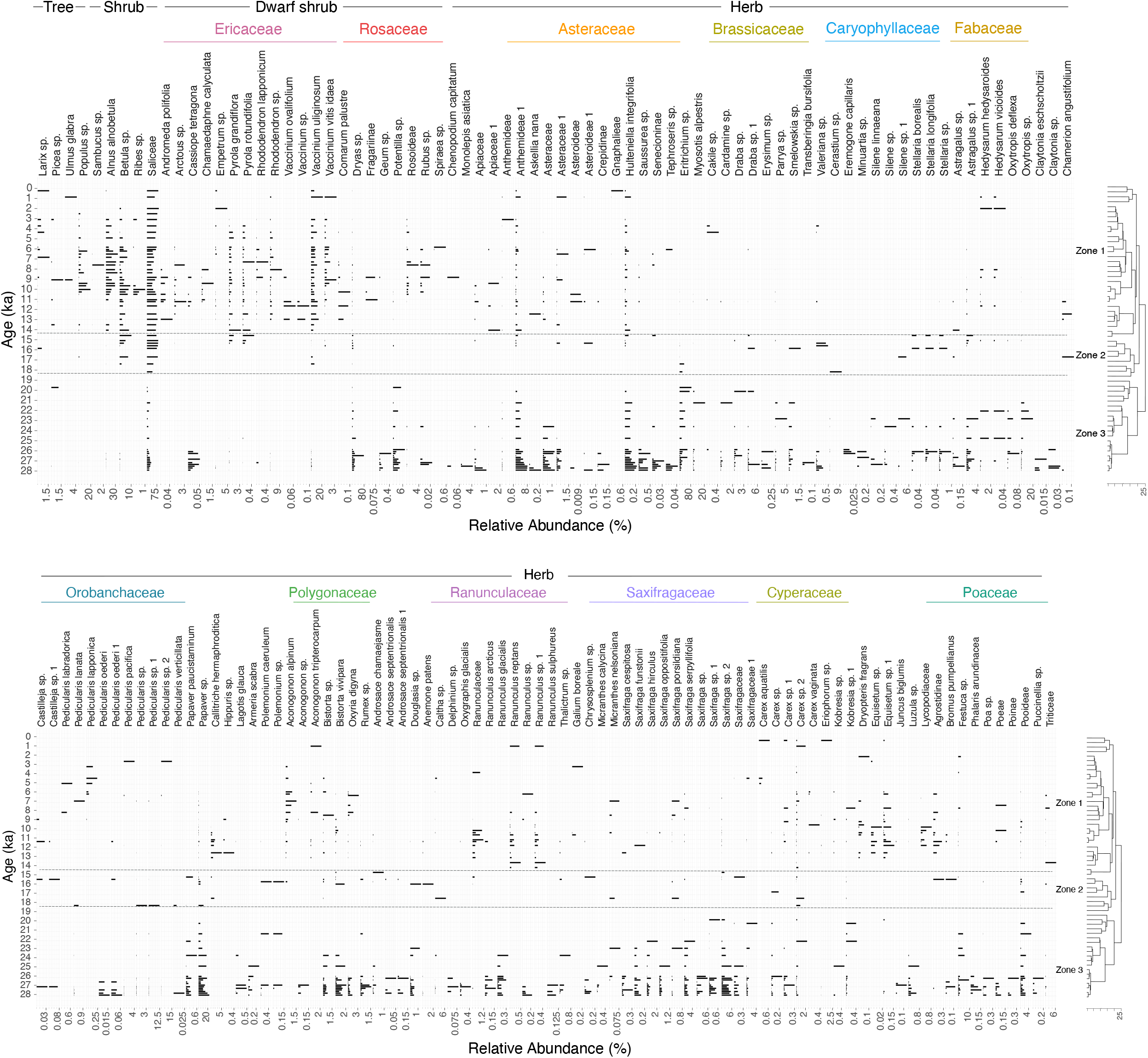

The PCA biplot (Fig. 3) of the first and second PC axes explain, respectively, 40.0% and 8.0% of the variance in the dataset. The ordination shows a clear separation of samples older and younger than 14 ka. Samples from 28–14 ka are mostly dispersed toward the negative end of the PC1 axis, with the highest loading on PC1 given by herbaceous *Eritrichium* sp. and *Papaver* sp. Along the positive PC1 axis, samples since 14 ka form a cluster, characterised by high values of *Alnus alnobetula* and *Vaccinium uliginosum.*

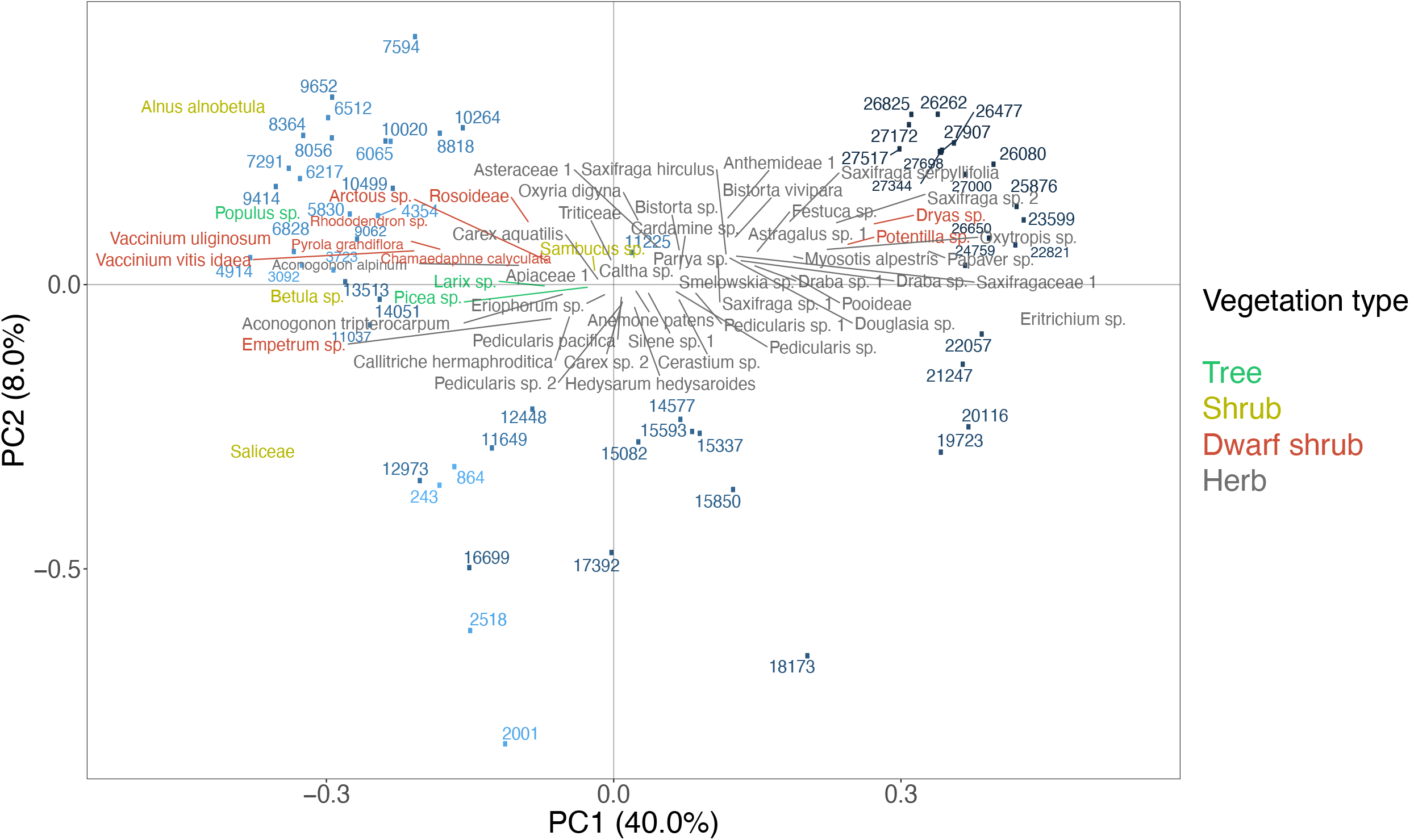

### Taxonomic alpha and beta diversity

We infer the highest species richness in the plant communities from 28–23 ka and a second richness peak during 13–9 ka (Fig. 4). Overall, we find almost no relationship between richness and the number of reads (r = −0.26, p = 0.05), however, lowest read numbers post-LGM co-occur with the lowest richness. Similar to richness, the diversity indices and evenness also have high values before 21 ka (Fig. 4), but the second peak of these indices lags the second richness peak by 3000 years (9–6 ka).

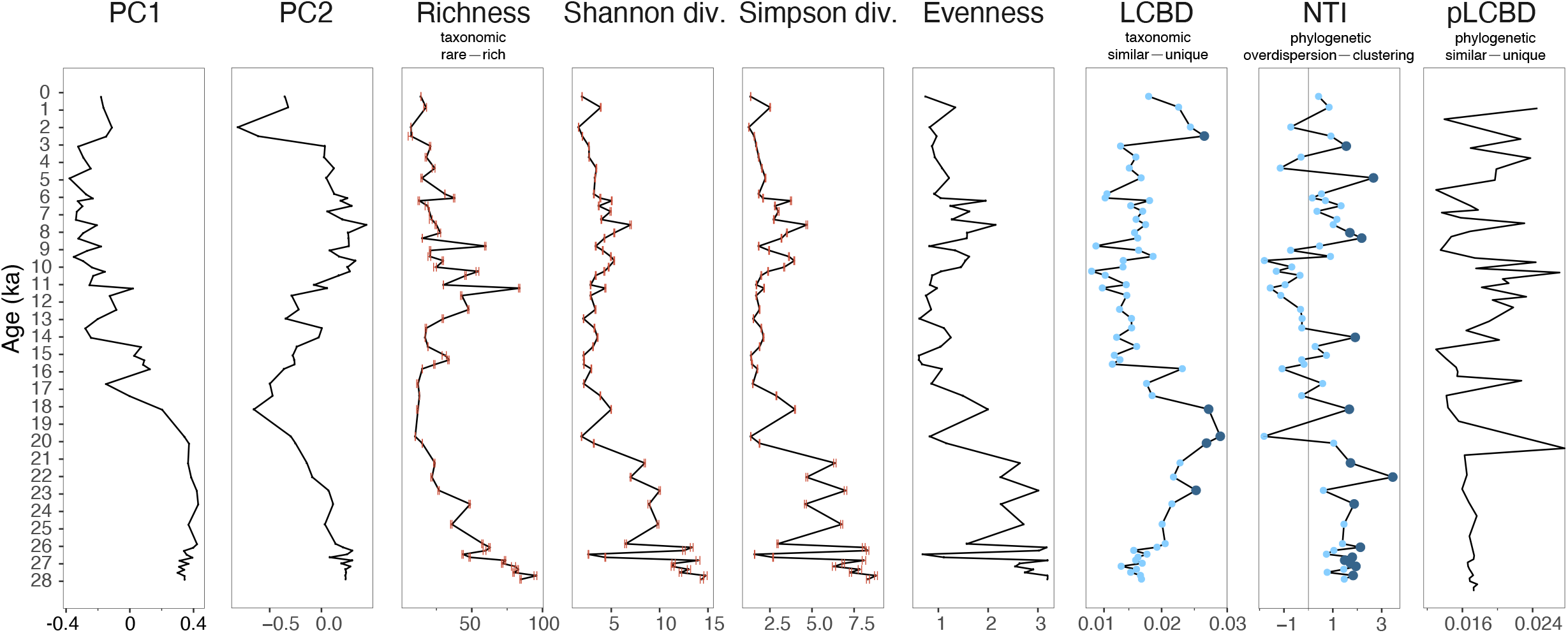

The abundance-weighted LCBD (Fig. 4) indicate that the plant populations are taxonomically unique during 24–18 ka as well as 3–2 ka. *Eritrichium* sp., Saliceae, and *Alnus alnobetula* contribute most to beta diversity (Supplementary Material Appendix 4 Fig. S2a). In contrast, the occurrence-based LCBD (Supplementary Material Appendix 4 Fig. S1) indicate significant uniqueness during 28–24 ka and 12–9 ka, i.e. times of high richness.

### Phylogenetic alpha and beta diversity

The phylogenetic tree (Fig. 5) enables an estimation of phylogenetic diversity indices. The abundance-(Fig. 4) and occurrence-based NTI as well as NRI (Supplementary Material Appendix 4 Fig. S1) show overall similar patterns. Both abundance- and occurrence-based NTI values are statistically significant and positive (p < 0.05) between 28 and 21 ka and between 9 and 5 ka, indicating a phylogenetic clustering in the plant community during these periods. The abundance-weighted NTI values (Fig. 4) during 28–14 ka (mean NTI: 1.06) are higher than those during 14–0 ka (mean NTI: 0.27), suggesting an overall increase in phylogenetic alpha diversity. Although not significant, the NTI shows markedly negative values during 14–10 ka (mean NTI: −0.47), indicating a tendency to phylogenetic evenness in the plant community.

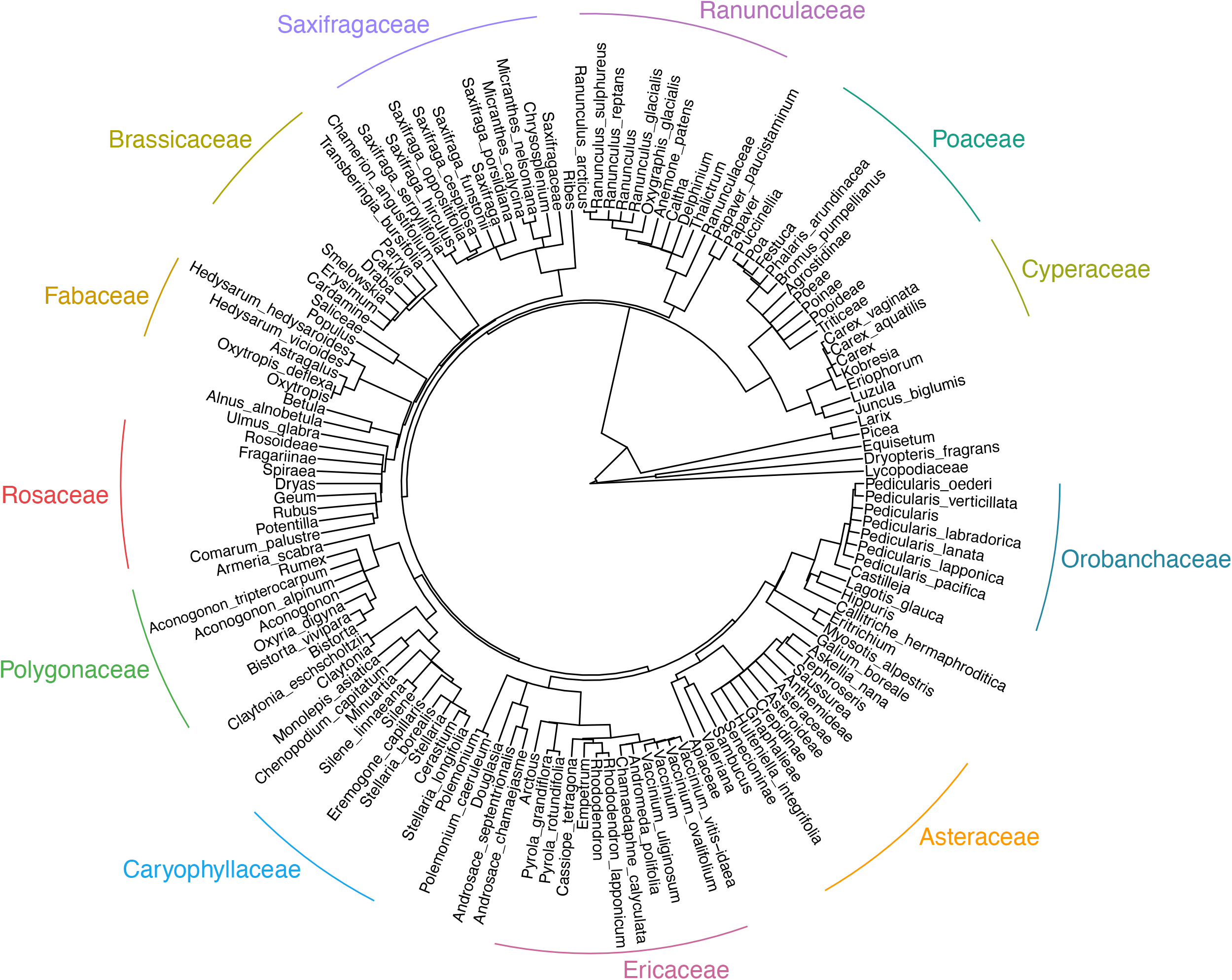

Both abundance-(Fig. 4) and occurrence-based (Supplementary Material Appendix 4 Fig. S1) pLCBD have higher values before 14 ka than later. This suggests that the latter period contributed more strongly to the phylogenetic beta diversity of the entire record.

### Relationship between taxonomic composition and phylogenetic diversity

The correlation of indices for phylogenetic alpha diversity, taxonomic composition, and phylogenetic beta diversity of the abundance- and occurrence-based plant communities are summarised in Table 1. In addition to the correlation over the last 28 k years, we report the results for the 28–14 ka and 14–0 ka periods.

**Table 1.**
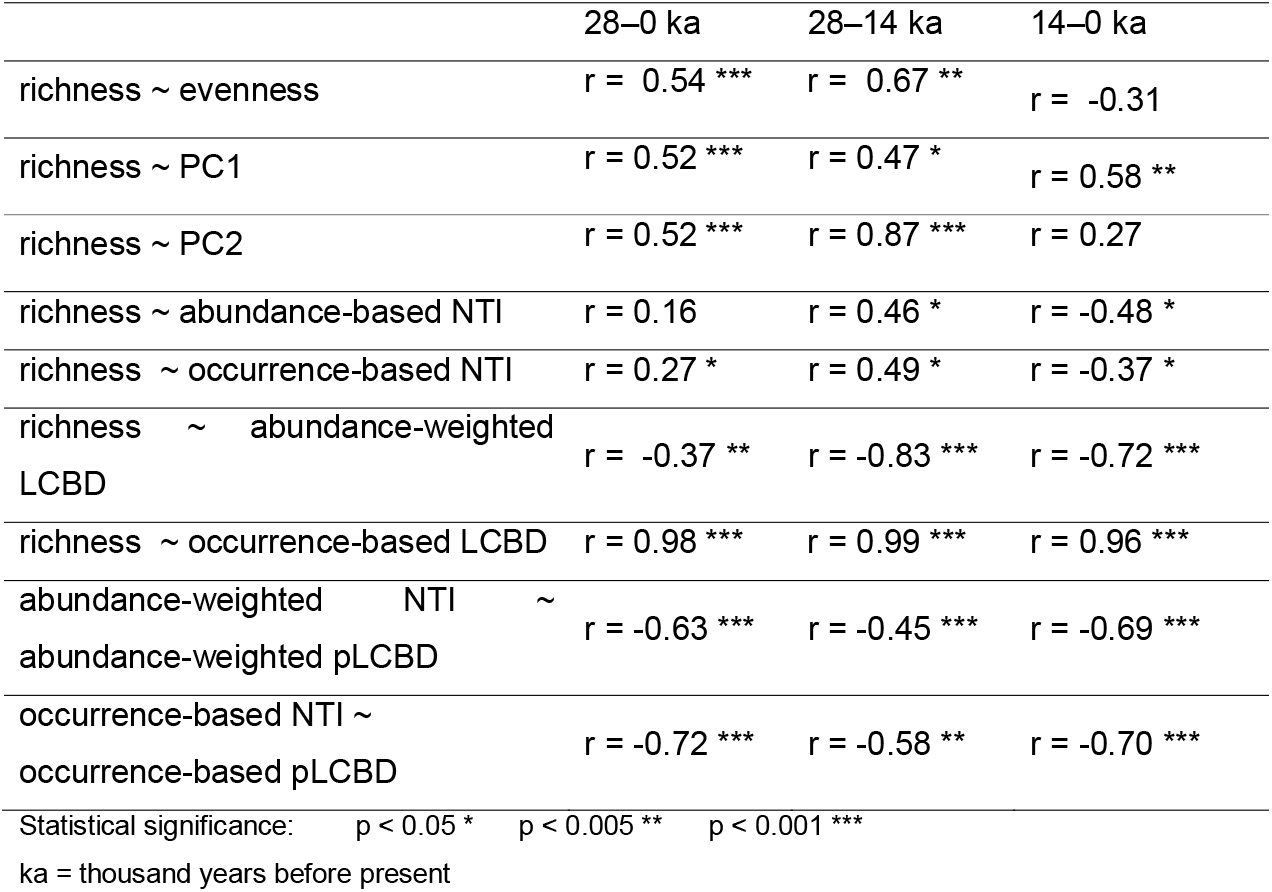
Pearson’s correlation between community composition and taxonomic and phylogenetic diversity in the plant communities inferred from sedimentary ancient DNA from subarctic Far East Russia

NTI values are significantly positively correlated with richness during 28–14 ka (r = 0.46, p <0.05, Table 1). In contrast, NTI values are negatively correlated with richness during 14–0 ka (r = −0.48, p < 0.05, Table 1). This indicates that rich communities are associated with closely related species during the cool period of 28–14 ka, but with more distantly related species afterwards.

Species richness is negatively related to the abundance-weighted LCBD (r = −0.37, p < 0.005, Table 1) for the whole record, whereas a strong positive relationship is found between species richness and occurrence-based LCBD (r = 0.98, p < 0.0001, Table 1). This suggests that the rich communities are taxonomically unique when considering only presence and absence data in the community. When the abundance-weight of the species are considered, rich communities are taxonomically even.

NTI is negatively associated with pLCBD in both abundance- and occurrence-based data since 28 ka (p < 0.001, Table 1), suggesting that the phylogenetically unique communities are more common within distantly related species.

## Discussion

### Vegetation history revealed by sedaDNA

Our results indicate that forb-dominated steppe-tundra rather poor in shrubs occurred from 28–19 ka and was followed by Saliceae-shrub tundra during 19–14 ka. Since 14 ka Ericaceae dwarf-shrubs expanded in the area and the lowlands in the Lake Ilirney catchment were covered by deciduous erect shrubs including *Betula* and *Alnus,* reaching a maximum density from 9–6 ka. Since the mid-Holocene, patches of open *Larix* forest expanded, several thousand years later than the maximum vegetation density and warming.

Zone 1 (28–19 ka) is characterised by forb-dominated steppe landscapes. Forbs such as Asteraceae, Saxifragaceae, *Dryas,* and *Eritrichium* (Boraginaceae) are common in the lake’s vicinity. This agrees with other ancient DNA-based studies in south-eastern Siberia (Courtin et al. submitted) and west Beringia of similar age (Willerslev et al. 2014). Our results suggest a dominant role for *Eritrichium* during the glacial period. As an alpine cushion nurse plant (Giguet-Covex et al. 2019), *Eritrichium* may have facilitated diversity under harsh environments through interactions with other species (Cavieres and Badano 2009). Shrubs such as *Betula*, *Alnus alnobetula*, Saliceae, and Ericaceae are absent or occur only at very low abundance. This contrasts to pollen records that show a relatively high percentage of shrubs from the same core (Andreev et al. submitted), which likely reflect a long-distant rather than a catchment-scale signal. As with previous sedaDNA evidence, we do not find any indication of a grass-dominated landscape as suggested by pollen records (Guthrie 2013). The taxonomic composition does not provide evidence that the formation of the forb-dominated landscape was mainly shaped by mammals (Geel et al. 2019).

Zone 2 (18–14 ka) is significantly different from Zone 1 in its community composition. Compared to Zone 1, forb taxa such as Asteraceae, Brassicaceae, and Cyperaceae greatly reduce their abundance, whereas dwarf birch and dwarf willow increase. This is probably related to climate amelioration during the Late Glacial period (Lozhkin et al. 2007). Increased amount of shrubs may also be related to a local signal of meltwater along the glacial channels (Whittaker 1993). The presence of *Betula* in the wider region is also confirmed from 18.7 ka by pollen records from the same core, though at low abundance (Andreev et al. submitted).

Zone 3 (14–0 ka) witnessed a shift from Saliceae-dominated dwarf-shrub tundra to erect-shrub tundra and open forest with Ericaceae understorey. High abundances of deciduous erect shrubs *(Alnus, Betula, Ribes)* and deciduous trees *(Populus)* occurred between 9 and 6 ka. This is also reported by other Siberian and Beringian pollen records (Szeicz and MacDonald 2001, Velichko et al. 2002, Anderson and Lozhkin 2015) and likely relates to increased summer warmth and moisture (Mann et al. 2002).

The period from 13–9 ka shows slightly reduced erect shrub values which might be related to cold events (Lozhkin 1993, Kokorowski et al. 2008). Interestingly the late Holocene is characterised by an invasion of *Larix,* although at low abundances. This implies *Larix* invaded the area after the Holocene Climatic Optimum; however this needs further investigation.

It appears that there is an overrepresentation of certain plant taxa in our sedaDNA record, for example, Saliceae, which is not dominant in the pollen records from Ilirney (Andreev et al. submitted). Willows preferentially grow along rivers and channels and hence have a higher chance of their vegetative remains being transported to the centre of a lake compared with upland taxa (Lozhkin et al. 2001). They invest massively in their roots and creeping stems, which increases their preservation potential (Pedersen et al. 2013).

By analogy to the past, future warming will result in increased summergreen shrub growth for upland areas replacing forb communities, which can already be observed via remote-sensing data of the last 20 years in the catchment. Furthermore, future changes in catchment hydrology will have a strong impact on the lowland vegetation, particularly willows. Our finding of a delayed forest response to warming implies that tree invasion strongly lags ongoing and future warming.

### Patterns of taxonomic alpha diversity and their relationship to community composition

The change in species richness shows that the greatest diversity of plant taxa occurred about 28 thousand years ago (Fig. 4), and was even higher than the maximum post-LGM richness, which occurred during 13–9 ka, and not, as expected, during the time of the Holocene vegetation “optimum” (as defined by the highest PC1 values) during 10–6 ka.

The specific geographical setting of our study area in central Chukotka probably favoured a higher biodiversity during the pre-LGM. As north-eastern Siberia was widely believed to be largely ice-free during most of the Pleistocene (Svendsen et al. 2004, Brubaker et al. 2005, Jakobsson et al. 2016), which when combined with the species-area relationship suggesting a positive association between area and species richness (Connor and McCoy 1979), then the enormous land area could have allowed more taxa to thrive during the pre-LGM and provided a glacial refugium for a variety of vegetation types during the LGM (Brubaker et al. 2005).

We find no evidence for the proposed mosaic of shrub and herb patches during the pre-LGM, which is a presumed characteristic of the mammoth steppe (Johnson 2009, Jørgensen et al. 2012). Hence, pre-LGM plant richness might not be related to the mammalian impact on vegetation (Sandom et al. 2014).

The LGM flora is largely a subset of pre-LGM flora. We find a decrease in richness compared to the pre-LGM, which is mainly led by the dramatic reduction of forb diversity, in particular with respect to Asteraceae, Brassicaceae, and Cyperaceae. This probably results from the local extinctions of thermophilic species in response to the LGM-related climatic deterioration (Stuart and LiSter 2007), or due to the potential breakdown of herbivore-vegetation interactions by declining mammalian populations during the LGM (Willerslev et al. 2014).

Interestingly, the pre-LGM forb diversity did not become re-established during the early Late Glacial. Either it was still too cold for many taxa, which is unlikely because the presence of Saliceae indicates a cool rather than very cold climate, or taxa invasion was delayed. The shift from herbaceous to woody plants during 18–14 ka may also have hindered or slowed the establishment of a species-rich forb community via an increase in shrub height and density altering the surface energy budget (Sturm et al. 2001) and changing the carbon balance and nutrient dynamics in the soil (Jackson et al. 2002). The second richness peak between 13–9 ka also coincided with a slight reduction in erect deciduous shrub abundance, while the abundance of dwarf shrubs (e.g. Ericaceae and Rosaceae) and herbs (e.g. Asteraceae and Ranunculaceae) increased.

Evenness, indicating a uniform distribution of many species, and richness are positively correlated for the 28–14 ka interval in our record. In contrast, the high evenness during 10–6 ka is accompanied by relatively low richness resulting in a weak negative correlation between evenness and richness during the last 14 kyr (Table 1). We assume that the densification of deciduous woodlands formed by *Betula, Alnus alnobetula,* and *Populus* for example, during the mid-Holocene may also have resulted in a disconnect between richness and evenness by inhibiting the establishment of a diverse herb and dwarf-shrub flora, resulting in the dominance of a rather species-poor undergrowth.

Several sources of bias may influence the diversity of the inferred vegetation. We used a short DNA marker in this study, which may limit the detection of families such as Cyperaceae and Poaceae due to their long sequence length (Sønstebø et al. 2010, Boessenkool et al. 2013, Alsos et al. 2018). The restricted specificity of the marker may have weakened the reliability of plant richness assessment in some families such as Asteraceae. Family-specific markers such as ITS primers (Willerslev et al. 2014) are needed for reducing the biases in chloroplast primers. Further alternatives to resolve the PCR biases is to use metagenomic approaches such as shotgun sequencing or capture approaches of enriching barcode regions (Gauthier et al. 2019).

Furthermore, sedaDNA diversity signals from Lake Ilirney may be impacted by changes in taphonomy (Giguet-Covex et al. 2019). For example, a large fluvial input and high sedimentation rate from glacier meltwater during the Late Glacial as a result of increased erosion of organic-poor sediments may have led to a dilution of DNA concentration and thus an apparent richness decline (Pawłowska et al. 2016). While the reference database we used for taxonomic assignment contains plants from the circum-arctic (Willerslev et al. 2014), not all species or even genera from our study area are included. The establishment of a reference database for Beringia is therefore a future task.

Our record indicates that a low shrub abundance may have enabled a rich forb-dominated steppe-tundra vegetation community during the pre-LGM while the dominance of deciduous erect shrubs during 10–6 ka coincides with low richness, but by an evenly distributed community. Our data imply that the vast expansion of shrubs in the currently warming Arctic (Chapin et al. 2005, Walker et al. 2006) may represent a regime shift for plant diversity. By analogy to the past, the ongoing “Arctic greening” will result in a reduction in plant richness in the long-term.

### Relationship between richness and phylogenetic alpha and beta diversity

Our exploration of the changes in phylogenetic diversity using plant sedimentary ancient DNA data reveals that species richness and phylogenetic clustering are positively related during the dry and cold 28–14 ka period but negatively during the wet and warm 14–0 ka period, indicating that rules of species assembly differed substantially during these periods.

The statistically significant and positive NTI values for 28–14 ka suggest phylogenetic clustering (Fig. 4, Table 1). Closely related taxa have more similar ecological preferences and often share similar environmental tolerances (Cavender-Bares et al. 2009, Mayfield and Levine 2010). Plant assembly during the pre-LGM consists of closely related sequences assigned to forb families such as Asteraceae and Saxifragaceae (Fig. 2). Phylogenetic clustering likely happened through selection of climatically pre-adapted species or even adaptation to cope with severe glacial conditions (Marx et al. 2017, Kubota et al. 2018).

A positive correlation between NTI and richness indicates that rich communities were composed of closely related species during 28–14 ka (Fig. 4, Table 1). A common assumption is that genetically closely related species tend to compete more intensely than their distantly related peers, resulting in reduced coexistence (Webb 2000, Cahill et al. 2008). However, competition also increases the opportunities for ecological facilitation through niche differentiation (Gioria and Osborne 2014). Some closely related arctic forbs have been found to interact positively (Kolář et al. 2013). For example, species of Asteraceae are able to coexist with their sister clades through stabilising niche differences (Godoy et al. 2014).

There is a tendency of an overall increase in phylogenetic distance during the last 14 kyr compared with the 28–14 ka interval (Fig. 4), indicating a shift from phylogenetic clustering before 14 ka to phylogenetic overdispersion afterwards. Climate amelioration after 14 ka supported the invasion of many taxa in Lake Ilirney catchment. However, species arrival depended not only on climate but also on, for example, the distance to the glacial refugia (Alsos et al. 2015). Species recruitment was therefore likely to be rather erratic and neither environmental nor biological filtering were shaping the community, resulting in a phylogenetically overdispersed community between 14 and 10 ka.

A negative correlation between NTI and richness, as found after 14 ka, indicates a pattern that rich communities are distantly related (Fig. 4, Table 1). Following the stress gradient hypothesis (Bertness and Callaway 1994), less stressful environments during 14–10 ka were characterised by erect-shrub tundra and open forest composed of distantly related species. Those species interact positively while biotic filtering is of minor importance. Such beneficial ecological facilitation may have supported maintenance of a high productivity between 14 and 10 ka (Zhang et al. 2016). Increased NTI values during 10–6 ka are accompanied by reduced richness. This is probably due to the unstable co-existence within closely related shrubs. After the massive expansion of shrubs and trees during the early Holocene, the shrub species tend to be kept from being too similar through competitive exclusion. Alternatively, the closely related shrubs can have negative effects on the diversity of the understorey by competition for light, water, and nutrients (Barbier et al. 2008). Further experiments are needed to validate whether competition or facilitation dominates when close relatives of arctic shrubs coexist.

Our results indicate that ecologically unique communities are rich when considering only presence and absence in the community (Fig. 4, Table 1). In contrast, when analysing abundance-weighted data (Supplementary Material Appendix 4 Fig. S1, Table 1), ecologically unique communities are species-poor. Species with high occupancy across the entire time series and high total abundance (e.g. *Eritrichium* and Saliceae) contribute most to the abundance-based beta diversity (Supplementary Material Appendix 4 Fig. S2a). Species of intermediate abundance contribute most to the occurrence–based beta diversity (Supplementary Material Appendix 4 Fig. S2b) as they show strong abundance variations (Heino and Grönroos 2016, Silva et al. 2018).

A general pattern of negative correlation between NTI and pLCBD between 28–0 ka is not surprising (Fig. 4, Table 1). Phylogenetically close species tend to share similar ecological requirements and are therefore often found in the same community (Mayfield and Levine 2010, Kamilar et al. 2014). These communities then exhibit similar phylogenetic patterns (Shooner et al. 2018) and become phylogenetically even less unique.

The changes in phylogenetic diversity before and after 14 ka differ from those shown by taxonomic diversity, suggesting that inferred community assembly rules can be hampered if the analyses are based solely on taxonomic or solely on phylogenetic diversity (Chai et al. 2016). Using a combination of taxonomic data and phylogenetic data gives a more complete representation of community structure. With respect to ongoing warming, our results imply, that, on the short-term, many species may invade the area leading to rich phylogenetic overdispersed communities. However, expansion of shrubs might cause a reduction in plant taxonomic richness phylogenetic clustering.

## Declarations

### Author contribution

Sichao Huang: conceptualization (lead); formal analysis (lead); writing - original draft (lead); writing – review & editing (lead). Kathleen R. Stoof-Leichsenring: conceptualization (supporting); formal analysis (supporting); resources (supporting); writing – review and editing (supporting). Sisi Liu: writing – review and editing (supporting). Jeremy Courtin: writing – review and editing (supporting). Andrei A. Andreev: writing – review and editing (supporting). Luidmila. A. Pestryakova: resources (supporting); writing – review and editing (supporting). Ulrike Herzschuh: conceptualization (lead); resources (lead); writing – review and editing (lead); supervision (lead).

