## Supplementary Material for "Plant sedimentary ancient DNA from Far East Russia covering the last 28 ka reveals different assembly rules in cold and warm climates"

### Appendix 1

#### Study area

As an area with unique vegetation composition that harbours many endemic species (Kuzmina et al. 2011), Chukotka is a hotspot for biodiversity in the Russian Arctic. It is a key region for understanding both Quaternary glaciation history and transcontinental species migrations as it lies at the far eastern edge of Asia bordering the Bering Sea (Kuzmina et al. 2011), which is known for being a glacial refugium during the last glacial maximum (LGM) (Brubaker et al. 2005). Chukotka is characterised by a comparatively pristine environment including wild herbivore mammals which have survived human impact.

#### Fieldwork

Sediment core “16-KP-01-L02 Long 3” was obtained from Lake Ilirney (67.34148, 168.30443) in summer 2016 as part of a joint Russian-German Expedition. The coring was accomplished using a UWITEC gravity corer equipped with a hammer action (Vyse et al. 2020). The 235 cm-long sediment core was cut into 1-m-long pieces and transferred to the Alfred Wegener Institute for Polar and Marine Research (AWI) in Potsdam, Germany where it was stored at 4°C prior to sub-sampling.

#### Subsampling

The core segments were cut into two halves. One was stored as an archive at 4°C and the other was opened for subsampling. The core subsampling was performed in the climate chamber of the Helmholtz Centre Potsdam - German Research Centre for Geosciences (GFZ). The subsampling environment was under strict hygienic rules to prevent contamination with modern DNA. The surface in the climate chamber was cleaned with DNA Exitus Plus ™ (VWR, Germany) and purified water. All sampling tools were cleaned regularly when taking each sample (Champlot et al. 2010). Sediment at either end of each core section were removed as they were in contact with the plastic tube and considered unsterile (Parducci et al. 2017). DNA samples were taken every two centimetres with 5 ml disposable syringes, in which the anterior caps were previously cut off with sterile scalpels. In total, 58 DNA samples were collected into 8ml sterile tubes (Sarstedt) across the entire core. The collected samples were then stored at -20°C.

### Appendix 2

DNA isolation and polymerase chain reaction (PCR) setup were performed in the palaeogenetic laboratory of AWI. This lab is dedicated to ancient DNA environment and is located in a building devoid of any modern molecular genetic work. For hygiene maintenance, the lab is equipped with an antechamber and separate rooms and individual UV-working station for DNA extraction and PCR set-up. Generally, the lab is cleaned by nightly UV-irradiation with ceiling UV-lamps.

#### DNA extraction

The 58 samples selected for DNA extraction range from 1 to 235 cm depth. DNA isolation was performed using a PowerSoil™ DNA Isolation Kit (MoBio Laboratories, California). The DNA extraction was divided into 10 batches. To check for chemical contamination, one extraction blank was included for each extraction batch. Approximately 3 g of sediment sample, 15 mL bead solution, 1.2 mL C1 buffer, 400 µL of 2 mg L^–1^ proteinase K (VWR International) and 100 µL of 5 M dithiothreitol were prepared in a bead tube for each sample. The samples were incubated at 56°C in a rocking shaker overnight. The subsequent steps were carried out according to the instructions of the manufacturer by Qiagen. In the final elution step, 800 µL elution buffer (C6) was added to the filter membrane and the incubation time was extended to 10 min. This step was performed twice and the final volume of the isolated DNA was 1.6 mL. Isolated DNA was stored at -20°C.

#### Polymerase chain reaction (PCR)

PCR protocols were set up with plant-specific *g* and *h* primers of the *trn*L gene (Taberlet et al. 2007). To distinguish the samples after sequencing, both forward and reverse primers were modified on the 5’ end by adding eight unique randomised nucleotides (Binladen et al. 2007). The primers were elongated by three additional unidentified bases (NNN) for improving cluster detection on Illumina sequencing platforms (De Barba et al. 2014). The PCR setup contained 1.25U Platinum® Taq High Fidelity DNA Polymerase (Invitrogen, USA), 1× HiFi buffer (Invitrogen, USA), 2 mM MgSO_4_, 0.25 mM mixed dNTPs, 0.8 mg Bovine Serum Albumin (VWR, Germany) 0.2 mM of each primer and 3 µL of DNA template. Each reaction had a total volume of 25 µL. PCRs were run in the Post-PCR area and were performed in a TProfessional Basic Thermocycler (Biometra, Germany) with the conditions of initial denaturation at 94°C for 5 min, followed by 50 cycles at 94, 50, and 68°C each for 30 s and a final extension at 72°C for 10 min. The PCRs were carried out in three replicates using different primer tag combinations. Twenty-one PCR reactions were performed, and 9–11 sediment samples were included in each reaction. PCR no template control (NTC) and the extraction blank were run alongside each reaction. In total there were 216 PCR products including replicates, NTCs, and extraction blanks. The expected amplification was validated by 2% agarose (Carl Roth GmbH and Co. KG, Germany) gel electrophoresis.

#### Purification, pooling and sequencing

The PCR products were purified using the MinElute PCR Purification Kit (Qiagen, Germany), following the kit manufacturer’s instructions. The purified PCR products were eluted to a final volume of 40 μL. The DNA concentrations were quantified with the Qubit® dsDNA BR Assay Kit (Invitrogen, USA) using 1 μL of the purified DNA. All samples were then pooled equimolarly to a final concentration of approximately 60 ng in 1711.80 μL. NTCs and extractions blanks were standardised to a volume of 10 μL and were also included in the pool. Library preparations and parallel high-throughput paired-end (2 × 125 bp) amplicon sequencing were performed on the Illumina HiSeq 2500 platform (Illumina Inc.) conducted by Fasteris SA sequencing service (Switzerland).

### Appendix 3

#### Processing the raw sequence data

Sequence data were processed using the OBITools package (Boyer et al. 2016). The raw forward and reverse reads were assembled to a single file using *illuminapairedend*. Based on an exact match to the tag combinations, the sequences were assigned to samples using *ngsfilter*. Identical sequences were then deduplicated using *obiuniq*. Sequences with less than 10 read counts were removed using *obigrep* to reduce possible artefacts. As a final denoising step, *obiclean* was used to exclude further sequence variants that likely originate from PCR and sequencing errors (Boyer et al. 2016). The reference used for taxonomic assignment is based on the curated Arctic and Boreal vascular plant and bryophytes database. It contains 1664 vascular plants and 486 bryophyte species (Willerslev et al. 2014, Soininen et al. 2015). The processed sequences were assigned using *ecotag* by searching for possible matches within the reference library.

#### Further sequence data filtering

To have a reliable taxonomic assignment of the sequences but not to overestimate the diversity, the data were further filtered. Only sequences assigned with a best identity of 100% and at family level or lower were kept. Rare sequences occurring with less than 10 read counts across the dataset were replaced with 0 as probable artefacts. Sequences that occurred less than three times in all the PCR batches including replicates were also discarded. After excluding extraction blanks and NTCs, the PCR replicates were summed up. Extraction blanks and NTCs accounted for 11,416,168 reads (25% of the total dataset). The contamination was most likely caused by the reagents in the PCR reaction and the bioinformatic pipeline. However, we kept the data from the two contaminated batches, as we identified the counts in the rest of the samples from these batches that have similar patterns as their corresponding replicates. Finally, we excluded sequences which were assigned to aquatics and bryophytes and kept only the terrestrials for this study. The raw sequencing data, related scripts for analysing and final sequence dataset with abundances and taxonomic assignments, are available in Dryad (*DOI link will be provided after the acceptance of the manuscript*).

### Appendix 4

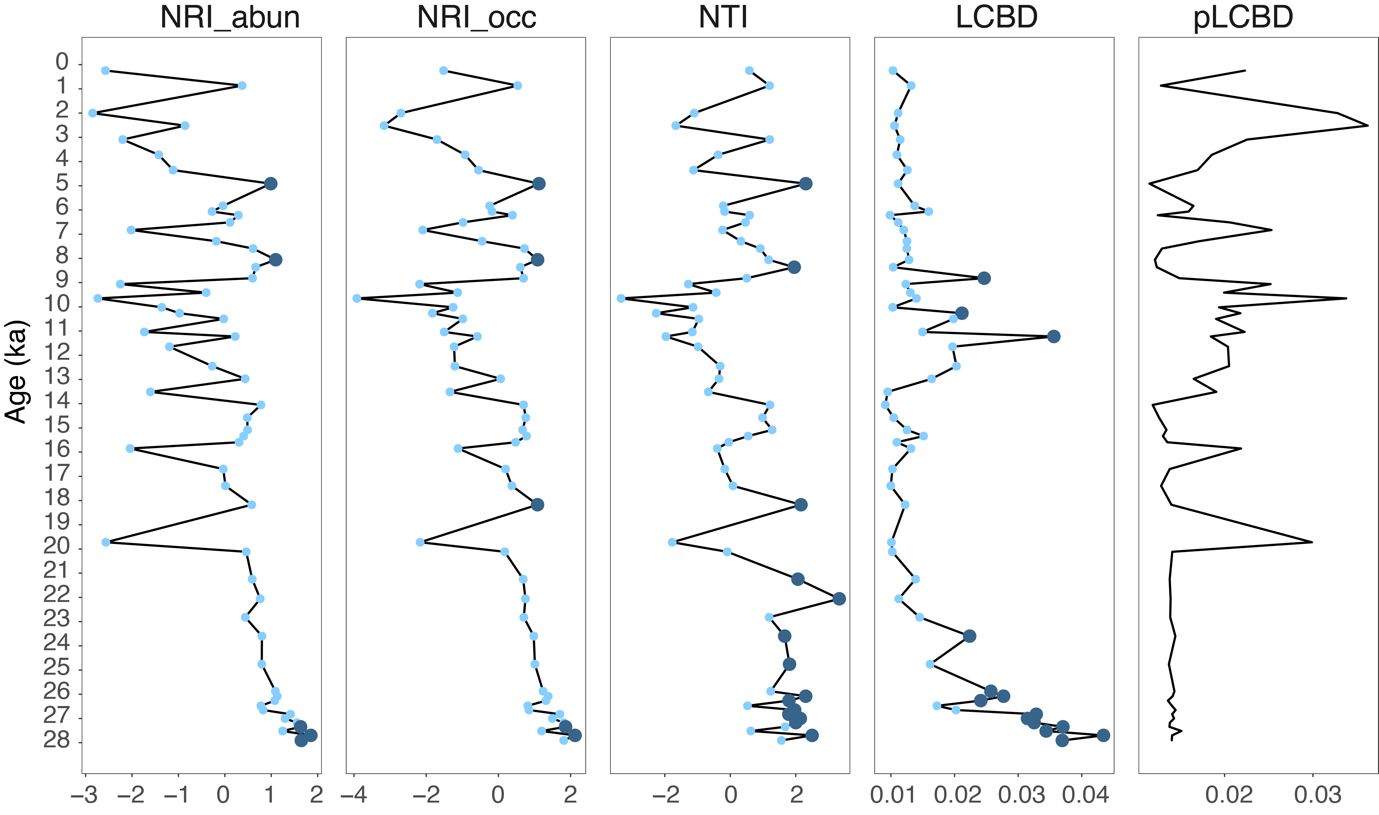

Fig. S1 Taxonomic and phylogenetic diversity in the plant communities from sedimentary ancient DNA. The measurements from left to right are abundance-weighted net relatedness index (NRI_abun), occurrence-based net relatedness index (NRI_occ), occurrence-based nearest taxon index (NTI), occurrence-based local contributions to beta diversity (LCBD), and phylogenetic occurrence-based local contributions to beta diversity (pLCBD). NRI, NTI, and LCBD values are coloured in dark blue when they are statistically significant (p < 0.05) and light blue when they are not significant (p > 0.05)

a

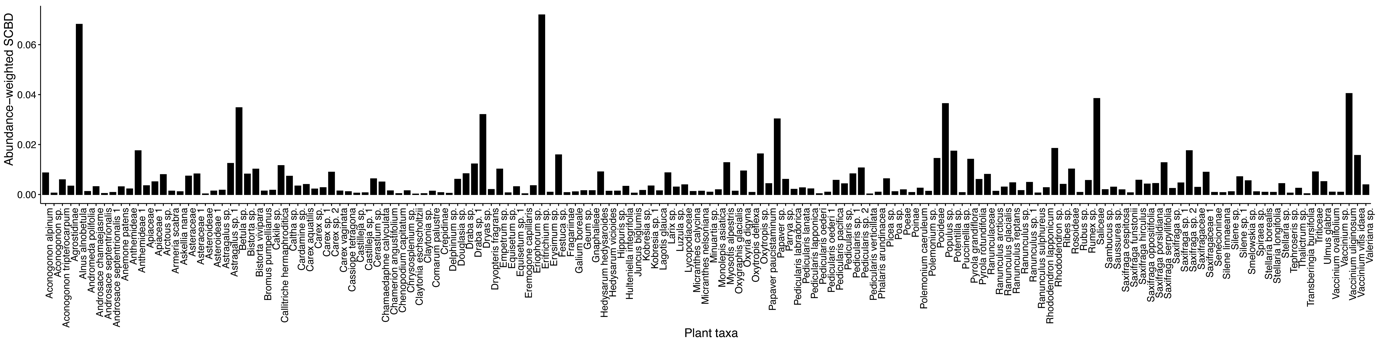

b

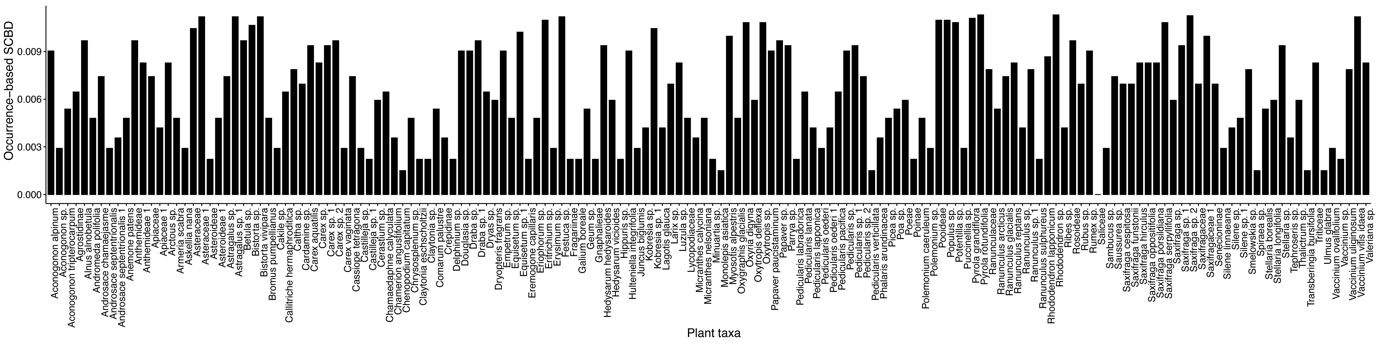

Fig. S2 Abundance-weighted (a) and occurrence-based (b) species contributions to beta diversity (SCBD) in the plant communities of sedimentary ancient DNA from subarctic Far East Russia
